# Global genetic rewiring during compensatory evolution in the yeast polarity network

**DOI:** 10.1101/2024.02.15.580535

**Authors:** Enzo Kingma, Liedewij Laan

## Abstract

Functional defects resulting from deleterious mutations can often be restored during evolution by compensatory mutations elsewhere in the genome. Importantly, this process can generate the genetic diversity seen in networks regulating the same biological function in different species. How the options for compensatory evolution depend on the molecular interactions underlying these functions is currently unclear. In this study, we investigate how gene deletions compensating for a defect in the polarity pathway of *Saccharomyces cerevisiae* impact the fitness landscape. Using a transposon mutagenesis screen, we demonstrate that gene fitness has changed on a genome-wide scale in the compensated strain. An analysis of the functional associations between the affected genes reveals that compensation impacts cellular processes that have no clear connection to cell polarity. Moreover, genes belonging to the same process tend to show the same direction of gene fitness change, indicating that compensation rewires the fitness contribution of cellular processes rather than of individual genes. In conclusion, our results strongly suggest that functional overlap between modules and the interconnectedness of the molecular interaction network play major roles in mediating compensatory evolution.

## 1. Introduction

Cells must perform a series of complex tasks to stay alive and replicate. The successful execution of these tasks depends on a dense network of proteins, DNA, RNA and other small molecules that physically interact. For various cellular functions, research in model organisms has unraveled the order and timescale of these interactions. Important examples of elucidated pathways are the endocytocic (1, 2), cytokinesis (3) and cell polarity pathways (4). Results obtained from model organisms are often extrapolated to other biological systems based on the assumption that the molecular mechanisms underlying the same function are shared among different species. While molecular conservation has been demonstrated in several cases, it is becoming increasingly clear that components essential for a function in one species may be absent in another species that performs the same function (5–8). Thus, pathway composition is flexible during evolution, even when the same biological functionality is retained.

One of the possible mechanisms that can drive the emergence of different pathways performing the same cellular function is compensatory evolution. During compensatory evolution, the fitness effects of deleterious mutations, often in the form of gene loss or loss-of-function mutations, are alleviated by the acquisition of secondary mutations elsewhere in the genome. These secondary mutations effectively restore the perturbed pathway without reverting the original defect, thereby exploiting degenerate mechanisms to perform the same function. How the architecture of the molecular interaction network within the cell provides opportunities for genetic compensation of functional defects is unclear (9, 10).

A prominent view is that molecular interaction networks are organized into dynamic modules (11, 12). Here, a module regulates a single physiological function and consists of a group of proteins that interact extensively with each other, but sparsely with the rest of the network. Proteins may belong to different modules at different points in time, making module composition dynamic (12, 13). Importantly, the modular organization of the interaction network has been reasoned to improve cellular evolvability (11, 14). Constructing a larger network from smaller, modular, building blocks that interact sparsely would limit the pleiotropic effects of mutations, allowing each function to be optimized independently during evolution. In line with this role for modularity during adaptation, laboratory evolution experiments have shown that compensatory mutations preferentially arise in genes that are functionally related to the gene that was initially perturbed (15, 16). However, for modules to truly evolve independently, compensatory mutations must not alter the genotype-fitness relationship of other modules. Whether the effects of compensatory mutations on the genotype-fitness landscape are indeed only local is currently unexplored. In addition, doubts have arisen regarding the relevance and benefits of modules in network evolution due to findings from models, computer simulations, and phylogenetic analyses (17–19).

An appealing model system to study compensatory evolution is the comparatively well-studied cell polarization pathway (20–22). Cell polarization remains functionally conserved across the diverse branches of the tree of life (23–25). However, components essential for polarization in one species can lack orthologs in other species (5), suggesting that compensatory evolution occurs naturally within this pathway. As in most eukaryotes, cell polarization in *Saccharomyces cerevisiae* is regulated by the cycling of the protein Cdc42 between a cytosolic inactive and a membrane-bound active state. Extensive research has uncovered the following three interconnected functional modules that collectively form the polarity pathway (26–29): actin-based transport, spatial cues and a reaction-diffusion module (figure 1a). While all three modules contribute to cell polarization, the reaction-diffusion module has been identified as its primary driver in wild-type cells. This functional module critically depends on the protein Bem1 (30, 31), which serves as a scaffold that connects most other components of the module. Deletion of Bem1 is highly disruptive and nearly abolishes the ability of cells to polarize (32, 33). Intriguingly, an earlier study has shown that this defect can be almost completely compensated by the additional deletion of two other proteins, Bem3 and Nrp1 (33). Bem3 is a known component of the reaction-diffusion module (34, 35), while Nrp1 is a protein of unknown function for which a role in polarity establishment might still be uncovered.

**Fig. 1.**
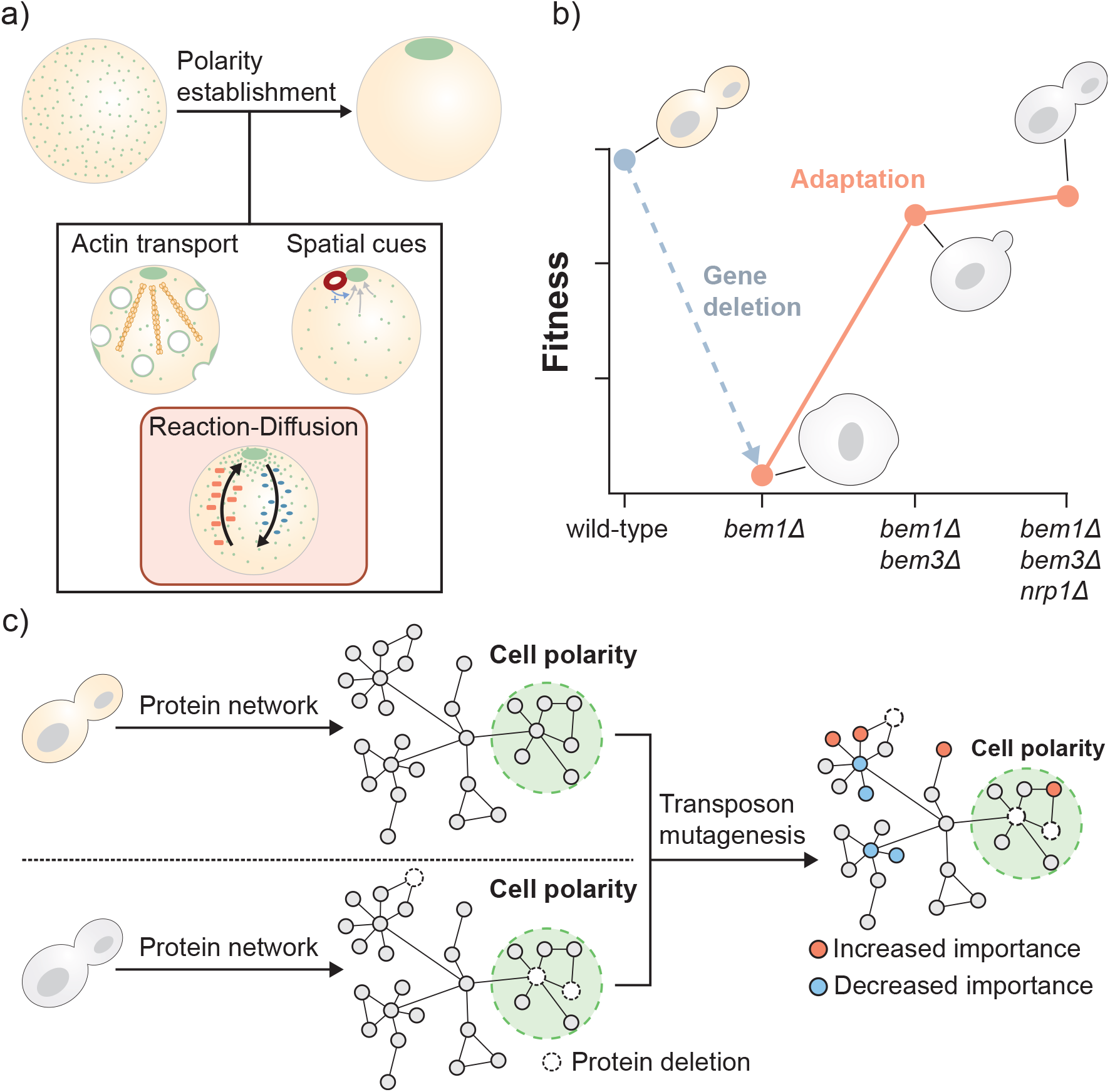
Determining the consequences of compensatory evolution of the polarity pathway on global cellular physiology. (**a**) Polarity establishment is an evolutionary conserved function of nearly all cell types. In budding yeast, cell polarization is initiated by the formation of an asymmetric distribution of Cdc42 (shown in green). Actin-based transport, spatial cues and a Bem1-mediated reaction-diffusion module have all been suggested to be involved in Cdc42 polarization dynamics. In this study, we examine how the fitness landscape changes after evolutionary compensation for a genetic perturbation in the reaction-diffusion module. (**b**) The phenotypic effects resulting from the deletion of the polarity protein Bem1 can be compensated by the additional loss of Bem3 and Nrp1. The resulting polarity mutant has a fitness similar to a wild-type strain under standard laboratory conditions. The figure is based on previous data obtained by Laan et al. (33). (**c**) We use a transposon mutagenesis screen to determine the effects of compensatory evolution on the genotype-fitness relationship on a global scale. Genes that differ in fitness between the wild-type strain and the polarity mutant are anticipated to encode proteins with modified physiological roles.

In this study, we evaluate how compensatory gene deletions within the cell polarity pathway influence the global fitness landscape and explore the role of modules in shaping evolutionary dynamics. We utilize a transposon mutagenesis screen to determine changes in gene fitness, defined as the impact of a gene deletion on fitness, and demonstrate that compensatory evolution affects the fitness landscape on a genome-wide scale. Despite compensation being achieved solely by gene deletions, the number of genes with increased and decreased fitness in the compensated strain are nearly equal. This finding indicates that evolution through gene loss does not always reduce mutational robustness. Because the evolutionary dynamics of proteins are often linked to their topological role in the protein interaction network (36, 37), we attempted to explain our observed pattern of gene fitness changes using the structure of the protein-protein interaction network of *S. cerevisiae*. However, we were unable to find any meaningful correlation with the protein network structure. Instead, studying the functional associations among the affected genes revealed that genes related to the same process tend to share the same direction of gene fitness change. Thus, although gene fitness changes do not remain isolated to the originally perturbed module, functionally related genes appear to respond similarly during adaptation. Cell polarity was amongst the biological processes enriched for genes with a negative fitness change, suggesting that redundancies between different modules within the polarity pathway mediate the capacity to compensate for gene loss. Collectively, our data demonstrates that mutations compensating for defects in a single functional module can lead to genome-wide changes in the gene-fitness relationship.

## 2. Results

### A. Compensatory mutations cause global changes in the fitness landscape

The unperturbed wild-type strain and the compensated *bem1* Δ*bem1* Δ*nrp1* Δ mutant, which we refer to as the polarity mutant, have similar fitness under laboratory conditions (33). However, whether the recovery of fitness after the loss of Bem1 is accompanied by restoration of the original fitness landscape has not been determined. The extent to which the fitness landscape is affected will reflect the molecular mechanism through which the gene deletions provide compensation. We considered the following two mechanisms for compensation to be likely.

The first mechanism relates to the argument that gene loss events during evolution often occur as a form of redundancy reduction (38). Similarly, modules can be redundant because they have largely overlapping functions. In the case of the polarity mutant, this implies that the reaction-diffusion module is redundant with one or more other modules that can drive polarity establishment. Deleting Bem1 causes the reaction-diffusion module to malfunction and interfere with these redundant modules. Consequently, further inactivation of the reaction-diffusion module through the deletion of *BEM3* and *NRP1* becomes advantageous. A recent study introduced a mathematical model that explains how functional redundancies between modules could be leveraged to form a latent pathway for polarity establishment (39). If compensation depends solely on the activation of these redundant mechanisms, we expect the fitness landscape to remain largely unchanged, with the exception of the decreased dispensability of the modules that are part of the latent pathway.

The second mechanism depends on the compensatory gene deletions facilitating novel interactions between other, non-mutated, proteins. According to the structure-function paradigm, a protein adopts a unique three-dimensional structure that ultimately determines its function. However, advancements in structural biology have revealed that protein structures are significantly more dynamic than previously believed (40). For example, up to 45% of residues in the eukaryotic proteome are found in intrinsically disordered regions of proteins (41), which lack a well-defined structure. Importantly, this structural flexibility enables protein function to adapt in response to varying cellular contexts and binding partner availability (42). Thus, the deletion of Bem3 and Nrp1 could compensate for the loss of Bem1 by modifying the functional properties of other proteins in the proteome, potentially leading to extensive rewiring of the interaction network. We anticipate that such rewiring of the protein-protein interaction network will result in global changes to the fitness landscape.

To uncover differences in gene fitness between the wild-type strain and the polarity mutant on a genome-wide scale, we performed the transposon mutagenesis screen SATAY (43). This screen utilizes the MiniDS transposon to generate a library of gene disruption mutants. In short, a plasmid carrying the MiniDS tranposon is transformed into the cell. By adding galactose to the growth media, the transposon is induced to migrate from the plasmid to a random location within the genome. This genomic integration disrupts the original sequence, typically abolishing gene function when it occurs within a coding region. By performing this process in a population of cells, a library of gene disruption mutants is created, with each mutant carrying a transposon at a different genomic location. The mutant library is subjected to a pooled fitness assay immediately after its generation. Subsequently, the genomic DNA is extracted from the mixed population and sequenced. Here, the genomic regions flanking the transposon insertion site serve as tags to distinguish the different mutants. The fitness of a mutant is reflected by the number of reads that align to the corresponding insertion site.

We generated six SATAY libraries of the wild-type strain and six libraries of the polarity mutant (figure 2a). Gene fitness was estimated by summing all read counts that map within a particular coding region. To compare datasets from independent experiments, the read count distributions were normalized (see section A) using a Beta-geometric normalization (figure S1), followed by a median of ratios normalization (figure S1). From the six replicates of each genetic background, we determined the mean (*µ*) and variance (*σ*) of the summed read read count for each gene (figure 2a).

**Fig. 2.**
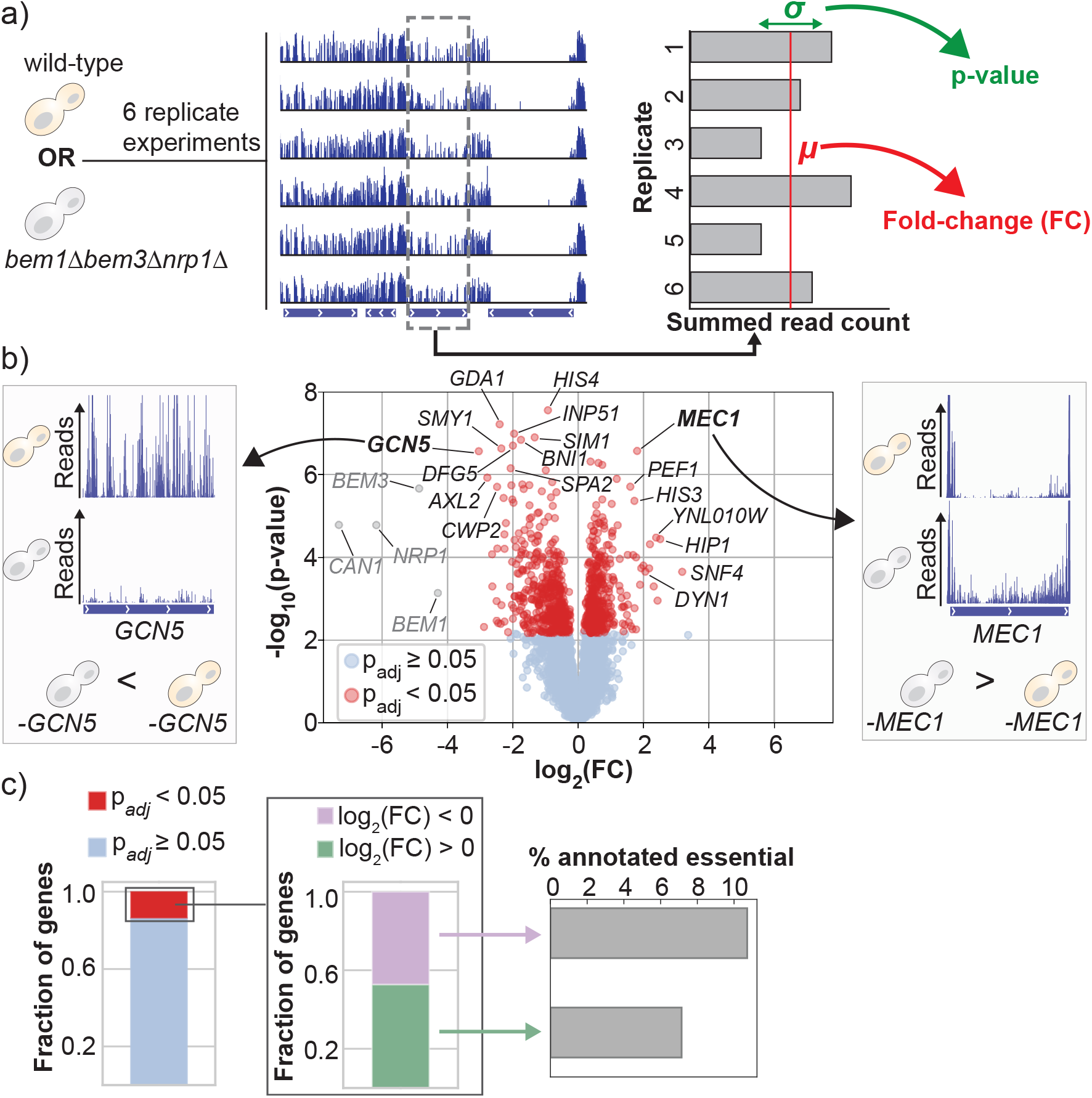
Compensatory gene deletions change the gene-fitness relationship on a genome-wide scale. (**a**) We created six replicate SATAY libraries of the wild-type strain and six replicate libraries of the polarity mutant. To estimate gene fitness, we summed the read counts of transposon insertions that map to the same coding region. The mean *µ* and variance *σ* of the total read count for each gene were calculated from six replicate libraries for each genetic background. The mean and variance were subsequently used to determine the log fold-change and significance of the difference in total read count between the two genetic backgrounds. (**b**) Volcano plot of the changes in gene fitness showing the log fold-change (FC) and the significance value of the differences in gene fitness between the two strains. Positive log fold-changes relate to genes that have a higher fitness in the polarity mutant relative to the wild-type strain. Statistical significance was determined with Welch’s t-test and corrected for multiple hypothesis testing with the Benjamini-Hochberg procedure. Genes that have been grayed out correspond to genes that were synthetically deleted from the genome of the polarity mutant during strain construction. (**c**) Approximately 13% of all annotated genes display a significantly altered tolerance to transposon disruptions between the two strains. An analysis of the sign of gene fitness changes in this set of genes shows that the ratio of genes that increase their relative fitness and those that decrease their relative fitness in the polarity mutant is nearly equal. Both the set of genes with increased and decreased fitness includes several genes that have been annotated as essential.

The changes in the mean total read counts between the wild-type strain and polarity mutant are presented in the form of a volcano plot in figure 2b. Interestingly, we find that a large fraction of the annotated genes in the genome have a significant difference in fitness between the wild-type strain and the polarity mutant. Specifically, a total of 883 genes, which constitutes more than 13% of all annotated genes in the yeast genome, has an altered fitness (figure 2c). As a reference, a comparison of SATAY libraries obtained from the same genetic background revealed no statistically significant differences (figure S2). When considering the sign of the changes in gene fitness, we find that the set of genes with increased fitness (414 genes) is nearly as large as the set of genes with decreased fitness (468 genes). Moreover, both sets include several genes that have been annotated as essential (figure 2c).

These data show that compensatory evolution affects a substantial fraction of the genome. Despite the intention to limit the initial perturbation to a single functional module (33), the effects of compensation on the fitness landscape extend far beyond this module. These widespread changes in gene fitness suggests that the recovery of cell polarity cannot be accomplished by merely reallocating the lost functions of the disrupted module to other redundant modules. However, while many of the genes with an altered fitness lack a clear relation to cell polarity, the set also includes several genes that are known to encode for polarity regulators (for example, *SPA2, BNI1* and *AXL2*). Functional redundancies between modules may therefore still mediate compensation, but seemingly not without affecting gene fitness regulating other cellular processes. Another surprising finding is that compensation through gene deletions causes a nearly equal number of genes to display increased and decreased fitness. Intuitively, restoring a pathway through gene deletions is expected to shift dependency to other cellular components that take over the lost functions. This indicates that the compensatory gene deletions reorganize the gene-fitness relationship rather than simply redistribute the lost functions to other, possibly redundant, genes.

### B. Changes in gene fitness cannot be predicted from the structure of the protein-protein interaction network

Are all genes equally likely to have a modified fitness in the compensated strain? If not, establishing a relationship between gene characteristics and features of the molecular interaction network could provide valuable insights for making evolutionary predictions. Earlier studies have indicated a correlation between a protein’s evolutionary rate and its position within the protein interaction network (37). For example, highly connected proteins (hubs) are more likely to be essential (44) and tend to evolve slower (36, 45) than less-connected proteins, possibly because they more frequently participate in important interactions (46, 47). These findings suggest that the topology of protein interaction networks provides relevant information to understand the evolutionary dynamics of individual proteins.

We constructed a protein-protein interaction network (PPI) to see if we could identify a correlation between the topology of this network and the gene fitness changes in the compensated strain. Protein interactions were taken from the BioGrid database (48, 49). Because the ability to derive a correlation between evolutionary dynamics and PPI topology depends on network quality, we only included interactions from the Multi-Validated dataset (see Methods). The resulting PPI network consists of 3799 proteins (nodes) and 17205 undirected interactions (edges). While the majority of the proteins are part of a single, fully connected, network, a small portion is disconnected from the main graph (figure 3a). In addition, the network represents only 65% of the estimated 5800 proteins in the yeast proteome and is therefore incomplete. For the remaining 35% of the proteome, the protein interaction pattern has not been validated by at least two independent studies or experimental methods. Among the absent proteins is Nrp1, which is one of the proteins that provides compensation for the loss of Bem1 when deleted. Despite these limitations, we successfully validated that proteins in our network with more interactions are more likely to be essential (figure 3b-c), a relation formally referred to as the centrality-lethality rule (44)). This demonstrates that crucial structural features of the network can still be inferred.

**Fig. 3.**
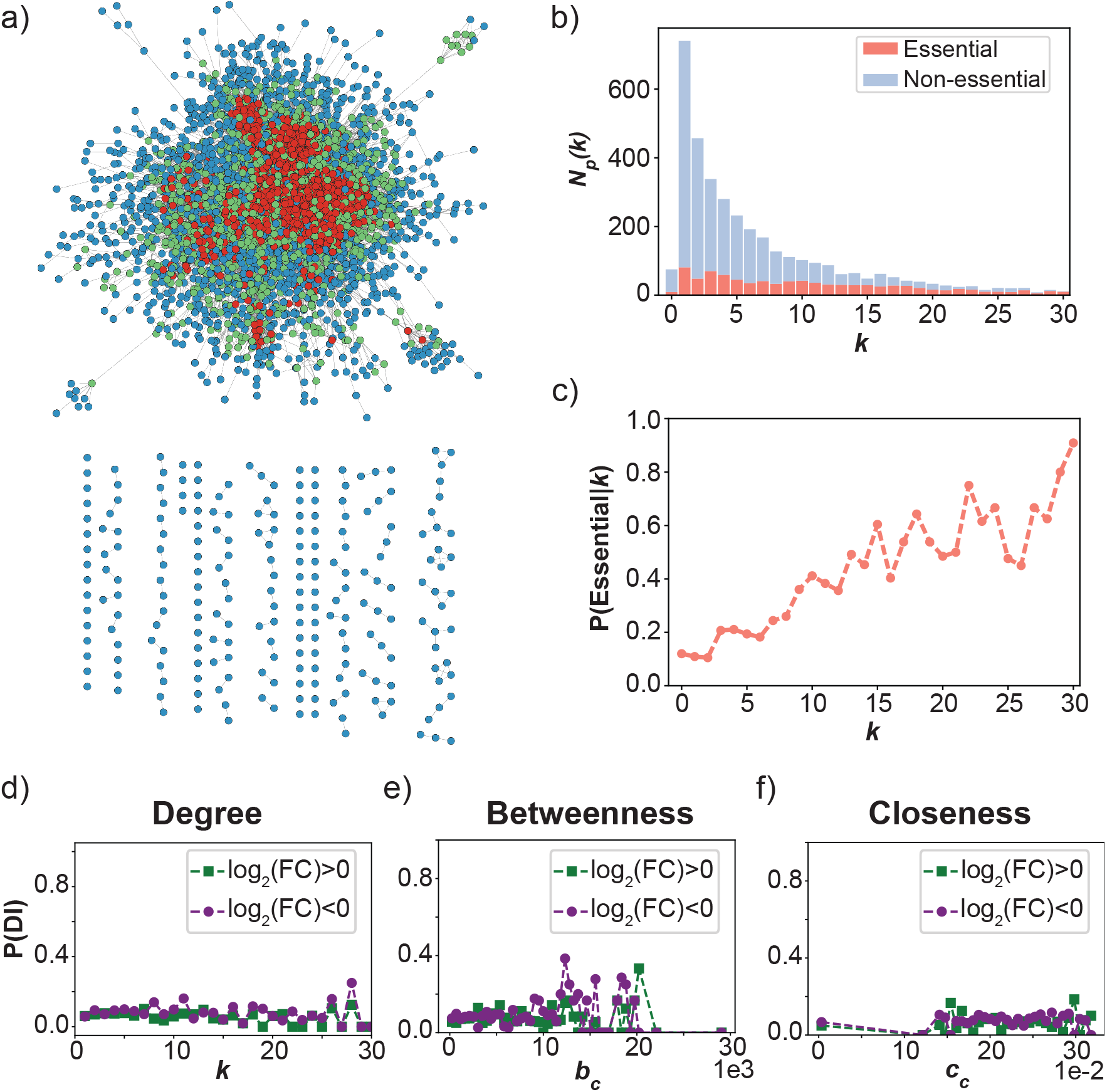
Likelihood of gene fitness change does not correlate with centralities of the protein-protein interaction network. (**a**) Visualization of the protein-protein interaction network constructed from the interactions annotated by the BioGRID database. Nodes are colored according to their degree *k* (blue: *k ≤* 3, green: 4 *≤ k≤*10, red: *k >* 10). (**b**) Stacked histogram of the number of proteins *N*_*p*_ (*k*) with degree *k*. Proteins for which the gene is annotated as essential are shown in red, non-essential proteins are shown in blue. The histogram shows that the distribution for *k* is roughly uniform for essential proteins, while it is skewed towards lower values of *k* for non-essential proteins. (**c**) The probability that a node is essential as a function of *k*. The graph displays an increasing trend for the probability that a node is essential for larger values of *k*. Combined with the histogram shown in (b), it can be seen that this increasing trend is due to an enrichment of essential proteins among those with a high degree, and not because most essential proteins have a high degree. (**d-f**) The conditional likelihoods for changes in gene fitness as a function of the degree (d), betweenness (e) and closeness (f) centralities of their corresponding protein in the network shown in (a). For the three centralities shown in panels (d-f), the likelihood distribution is approximately uniform.

The importance of a node for different topological features of a network can be described using network centralities (50). Here, we focus on three commonly used centralities: degree, betweenness and closeness. These centralities relate to the role of a protein in the PPI network. For example, hub proteins at the center of a module will typically have a high degree, while proteins that mediate connections between modules will have a high betweenness. We provide an overview of the distributions of these centralities in our network in figure S3b-d. Importantly, figure S3b shows that the degree distribution follows the sub-linear dependency characteristic of biological networks when plotted on a log-log scale. Furthermore, we show for 4 polarity proteins that those which strongly impact cellular viability (Cdc42 and Bem1) can clearly be distinguished from those which have a lesser effect (Bem3 and Bem2) based on their degree and betweenness (figure S3b and d). The relationship with closeness is, however, less clear (figure S3c).

Next, we attempted to derive a correlation between the three centrality measures described above and the likelihood that gene fitness has changed after compensatory evolution. However, the likelihood distributions are uniform for all centralities (figure 3d-f), indicating a lack of correlation. We therefore conclude that the changes in gene fitness are independent of the topology of the analyzed PPI network.

### C. Genes with an altered fitness are associated with a diverse array of cellular processes

The lack of a correlation between PPI network structure and the likelihood of observing a change in gene fitness prompted us to see if our dataset can perhaps be structured based on a different feature. Importantly, the lack of dependence on the PPI network does not exclude the possibility that genes participate in the same cellular process, as this does not require proteins to physically interact. These indirect functional relationships between genes can be captured with functional association networks.

We used the STRING database to create a functional association network from the genes with a differential fitness between the two genetic backgrounds (51–53). In this database, functional relations between genes are inferred by integrating information from multiple sources, such as text mining, molecular complex annotations and (predicted) physical or genetic interactions. Each interaction is scored based on quality of the evidence, allowing more weight to be given to high-confidence interactions.

Similar to our PPI network, the functional associations resulted in most genes being part of a single, connected, network from which only few nodes are disconnected. Analyzing the degree distribution indicated a scale-free structure of the network, suggesting it can be partitioned into smaller sub-graphs. We ran the Markov Cluster (MCL) algorithm (54) to uncover these sub-graphs, assigning interaction confidence scores as weights. This approach uncovered 120 clusters, of which 49 consisted of three or more genes. For the seven largest clusters, we performed a gene ontology (GO) enrichment test for biological process GO terms (figure 4b and figure S4). All seven clusters were significantly enriched for at least one GO term. Five clusters were enriched for the following terms related to cellular homeostasis: translation, metabolism, signalling, chromatin remodeling and ribosome biogenesis. The remaining two clusters were associated with components of the cell division machinery: cell polarity and microtubule dynamics.

**Fig. 4.**
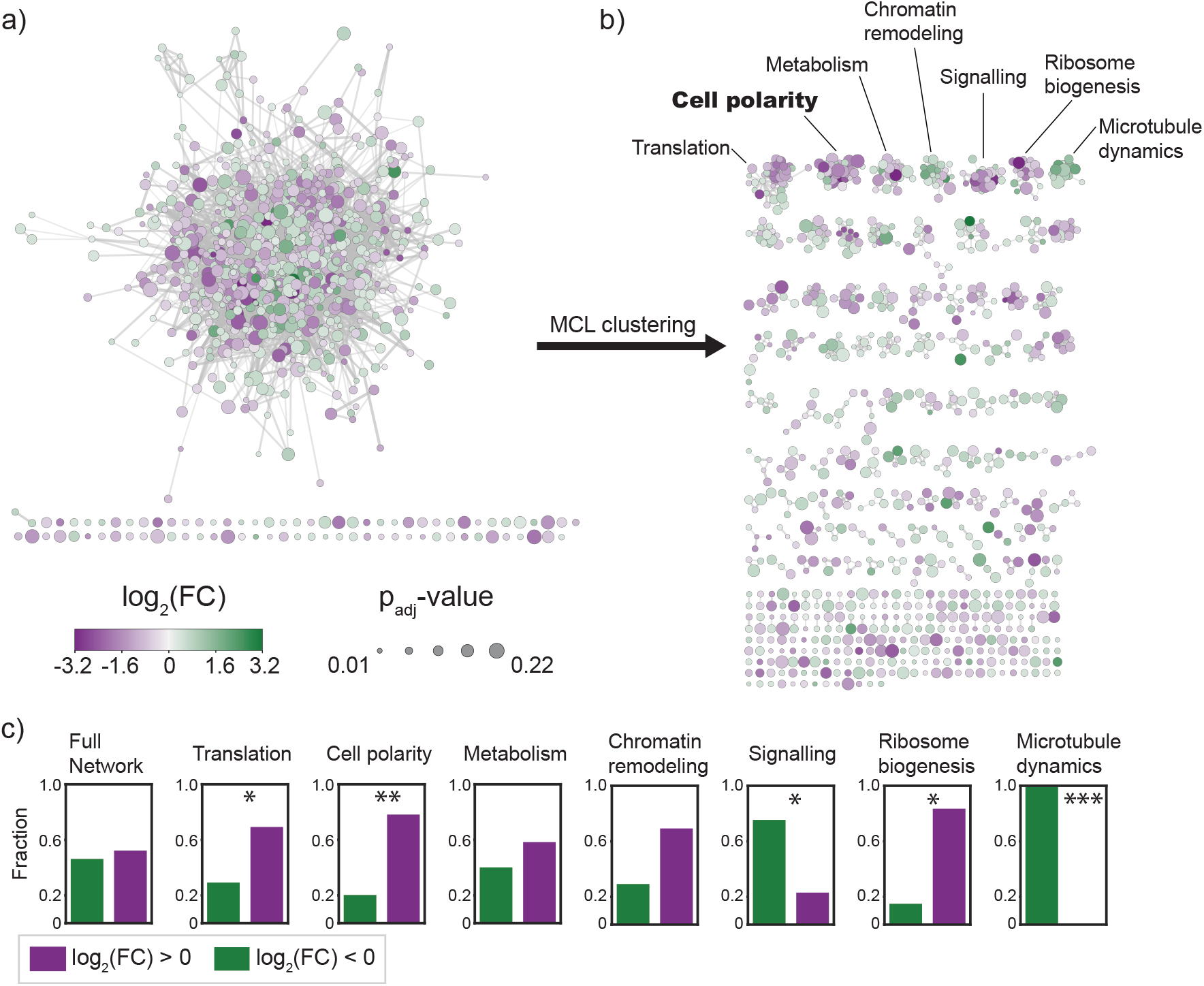
Clustering of a functional association network reveals diversity of biological processes affected by compensatory evolution. (**a**) Graph of the functional association network constructed from genes that have a differential fitness between the wild-type strain and the polarity mutant. Node color scales with the magnitude and sign of the fitness effect, node size scales according to the significance level (adjusted p-value). The graph shows that most genes are contained in a single, densely connected network. (**b**) The clusters formed from the graph shown in (a) by the Markov clustering algorithm (MCL). The clusters are arranged from large (top left) to small (bottom right). The biological process gene ontology enrichment is shown for the seven largest clusters. (**c**) Distribution of genes with a decreased (purple) and increased (green) fitness in the polarity mutant for the seven largest clusters. The clusters enriched for translation, cell polarity, and ribosome biogenesis are enriched for genes with decreased fitness. Conversely, the signalling and microtubule dynamics clusters are enriched for genes with increased fitness. *=*p <* 0.05, **=*p <* 0.005, ***=*p <* 5 *·* 10^*−*4^. The significance level was determined with Fisher’s exact test.

When assessing how genes with increased and decreased fitness are distributed among the clusters, we found that genes within the same cluster were often, but not always, more likely to share the same sign in fitness change than would be expected from a random partitioning (figure 4c). specifically, the clusters enriched for genes related to translation and cell polarity were also enriched for genes with an increased fitness. Alternatively, genes in the signalling and microtubule enriched clusters were more likely to have decreased in fitness.

Our analysis reveals widespread functional connections between the genes in our dataset. At the same time, by partitioning their functional association network into sub-graphs we were able to identify that these genes are involved in a diverse set of cellular processes. Intriguingly, the changes in fitness dependency seem to occur at the cellular process level rather than being attributed solely to individual genes. This uniform response of genes regulating the same process resembles the observed monochromatic behaviour of genetic interactions between modules of the yeast metabolic network (55). When comparing the cellular processes that are important for survival between the polarity mutant and wild-type strain, we observe several notable differences.

First, the general trend of genes involved in translation showing reduced fitness in the polarity mutant suggests that this strain is more sensitive to variations in protein copy numbers. This consequence of the compensatory mutations was also predicted by the mathematical model of Brauns et al. (39). Second, the finding that genes regulating microtubule dynamics become increasingly dispensable is surprising, given microtubules are required to perform vital cellular functions such as spindle positioning and nuclear migration (56, 57). Thus, compensatory evolution appears to profoundly affect the genetic wiring of the cell, changing the patterns of essentiality and dispensability beyond the genes related to the perturbed module.

### D. Compensatory evolution is mediated by redundancies within the polarity pathway

The wild-type strain and polarity mutant are nearly identical with respect to their polarization efficiency (33). The ability to polarize efficiently despite the loss of three genes implies the existence of an alternative pathway that is active in the absence of the reaction-diffusion module. Because it can function without adding any components, this pathway is considered to be latent or hidden within the interaction network of the wild-type strain (39). While reconstruction of the polarity mutant has demonstrated the existence of this latent pathway, its specific structure remains to be determined. In the previous section, we found a cluster enriched with polarity genes amongst those affected by the mutations compensating for defects in the reaction-diffusion module. The presence of this cluster implies that functional redundancies between the different modules of the polarity pathway could play an important role during compensatory evolution.

To uncover how these redundancies can be exploited to constitute an alternative polarization mechanism, we annotated the genes in the cell polarity cluster according to their gene ontology (figure 5a). This revealed that the majority of the genes encode for proteins that are part of one the following three cellular components: (1) cortical actin patches, (2) the bud scar and (3) the polarisome. Interestingly, each of these components relates to a different module of the known polarity pathway. Specifically, the bud scar provides a spatial cue that directs the axis of polarity, while cortical actin patches and the polarisome regulate the transport of active Cdc42 via endo- and exocytic vesicles, respectively. For all three cellular components, a substantial fraction (>20%) of the complete set of annotated genes is represented in our dataset.

**Fig. 5.**
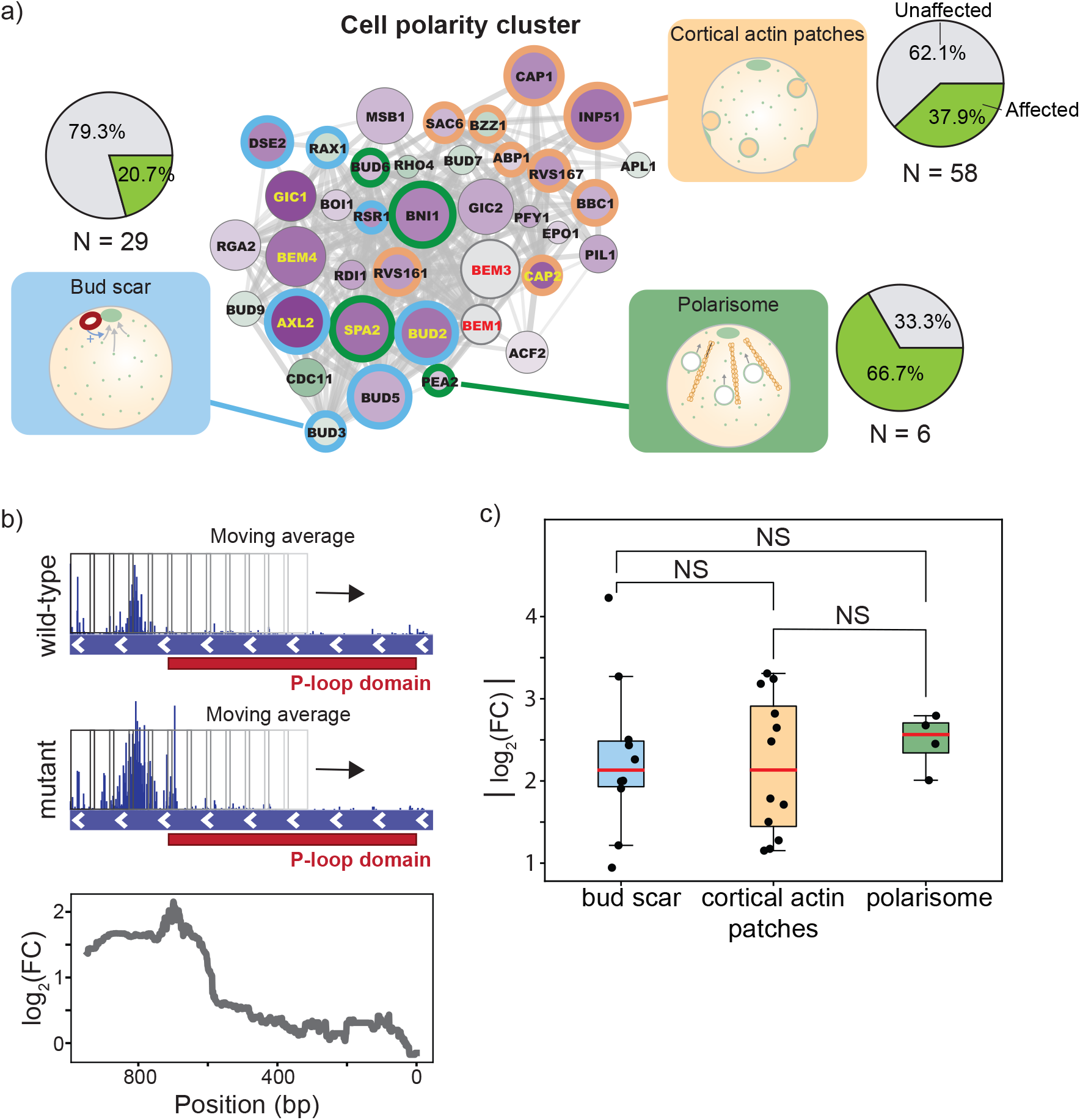
Redundant polarity modules contribute to the latent polarity mechanism in a non-hierarchical manner. (**a**) Genes in the cluster enriched cell polarity factors are annotated based on whether they belong to the bud scar (blue), cortical actin patches (yellow) or polarisome (green). The fraction of all *N* genes annotated with each of these three cellular component GO terms that are present in our dataset is shown in the pie charts on the right. For all three components, at least 20% of the annotated genes are present in our dataset. The cortical actin patch has the lowest representation, but is also the least polarity-specific of the three cellular components. (**b**) An example of the sliding average procedure taken to obtain the domain with the largest log-fold change is shown for *CDC11*. In *CDC11*, the region downstream of the P-loop domain (relative to the start codon) has an increased tolerance to transposon disruptions, while the remainder of the gene shows a similar tolerance between the two strains. Using this method, we can accurately capture this domain-dependency. The largest absolute value of the log-fold change profile over a gene was used to generate and compare the distributions shown in c. (**c**) Log-fold change distributions for the genes in our dataset encoding for proteins of the bud scar, cortical actin patches and polarisome. The orange line denotes the median of the distribution, the whiskers show the interquartile range. The three distributions all had very similar medians, and the difference in their mean was not statistically significant (p>0.05, t-test).

While all three cellular processes identified above are active in wild-type cells, their contribution to cell polarization remains inconspicuous when the Bem1-dependent reaction-diffusion module is active. This feature has been attributed to the hierarchical relationship that normally exists between modules in the polarity pathway (58). To determine if this hierarchy still exists within the latent pathway, we used the log-fold change distributions of genes belonging to each of the three modules. Specifically, the module that takes on the most prominent role in the latent pathway should show the greatest change in gene fitness.

However, quantitatively comparing log-fold changes in read counts summed on the gene level can give distorted results because their values depend on gene size. For example, if the region within a gene that exhibits a differential tolerance to transposon disruptions is small relative to the total gene size, the log-fold change value will be attenuated. To address this issue, we performed a more refined estimate of changes in gene fitness by using a moving average with a 300 bp window over the genes to obtain a fitness profile across the coding region (figure 5b). The choice for a window size of 300 bp was made based on the average size of a protein domain. Using this approach, we were able to identify the region within a gene that gives the largest log-fold change values between the wild-type strain and the polarity mutant. However, we note that while using a smaller window to compare genes allows a more accurate quantification of the log fold-change, it also increases the sensitivity to outliers.

When comparing the maximum log-fold changes in genes of the three modules described above, we observed no significant (p<0.05, t-test) changes between the means of their distributions (5c). Therefore, each module appears to be of equal importance for the functioning of the latent polarity pathway.

Although our results do not provide a complete molecular mechanism for the latent polarity pathway, they do offer important insights into its structure. The increased dependency on spatial cues we observe agrees with previous studies that have shown that the deletion of Bem1 is not lethal, but only if the bud scar remains intact (30, 32). This dependency on the bud scar therefore appears to persist even after mutations that compensate for the fitness effects caused by losing Bem1 have occurred. In contrast, the role of actin based transport in cell polarization has been a subject of ongoing debate (26, 59–63). Our results support the role of the actin network in promoting polarity establishment, at least within the latent pathway. However, the lack of hierarchy between the different modules in the latent pathway suggests that,while they contribute to polarity establishment, each individual module contribution is too weak to drive polarization.

## 3. Discussion

In this report, we examined the effect of mutations compensating for a perturbation targeting the polarity pathway on the genotype-fitness relation. Remarkably, our data show that two strains, which polarize with nearly identical efficiency and differ by only three genes, exhibit fitness differences in over 13% of the genes in the genome. This percentage is similar to what has been reported for the percentage of genes that display a background-dependent fitness effect between different natural isolates (64, 65). Given that the genomic divergence between natural isolates is generally larger than that between the two strains we studied, we anticipated finding a smaller number of genes with a background-dependent fitness effect. One explanation for the relatively high number of affected genes we identified is that our analysis method was aimed at detecting changes in gene fitness on a continuous scale, rather than categorizing genes simply as essential or non-essential. Our set therefore contains genes that would likely have remained undetected by other studies. However, despite the higher sensitivity of our method, the number of affected genes reported is likely still an underestimation. Importantly, because we sum read counts over the entire open reading frame, our analysis is relatively insensitive to domain effects on fitness when genes are large. Similarly, by only considering coding regions we are unable to identify changes in gene fitness that are related to changes in gene expression levels.

The impact of compensatory mutations on the fitness landscape may vary depending on the environment in which fitness is assessed. (66, 67). In a recent survey of changes in gene essentiality between natural isolates of *S. cerevisiae*, it was proposed that the majority of the observed variation was due to environmental factors (64). That is, it was argued that most genes that emerged as differentially essential in their screen were caused by strain-specific responses to growth in non-standard media. The mutagenesis screen we performed has a similar limitation. SATAY requires transposition to be induced in media in which galactose is the sole carbon source (43), while the phenotypic similarity between our two strains has only been verified for growth under standard conditions (33). We can therefore not exclude that we would have observed less changes in gene fitness if we had performed a screen in a different environment. However, we expect that the effect of the environment on our results is limited for the following two reasons. First, the overall genetic divergence between our strains should be smaller than what is typical for natural isolates that may have adopted different lifestyles to accommodate to their niche. A different study indeed showed that most genetic interactions are conserved across environments (68). Second, while we did identify a cluster enriched for metabolically related genes in our gene set, genes involved in galactose metabolism or respiration are not overrepresented in this cluster.

With the aim of making a first step towards predictive models, we assessed whether the genes that are affected by compensatory evolution can be identified based on their position in a PPI network (65). We did not observe any correlation between the likelihood that gene fitness changes and the degree, betweenness and connectedness centralities of the associated proteins. Thus, the genes that will have a differential fitness effect between the two strains cannot be inferred from these three centralities. While we were able to validate the presence of several features that are typically observed in PPI networks, we cannot reject the possibility that the lack of correlation is a consequence of poor quality of our PPI network. The accuracy of annotated physical interactions is know to substantially vary between datasets (69, 70). To exclude interactions with low confidence, we used the Multi-Validated dataset available from the BioGRID to construct our PPI network. However, network quality does not only depend on the confidence that an interaction exists, but also on the confidence that an interaction is absent (71). In addition, bias is a significant problem in the mapping of interaction networks. Weak and transient interactions are often discarded because they cannot be captured with high confidence by affinity based screens. These interactions could be of particular importance in cases where network structure needs to be recovered in response to gene loss, as recent work identified these interactions as the “glue that holds the cellular network together” (72).

The concept of modularity remains strongly embedded, both implicitly and explicitly, in cell and synthetic biology (16, 73). Using the functional associations available from the STRING database, we found that genes with differential fitness can be grouped into functionally enriched clusters, suggesting that some level of modular organization does exist within biological networks. This is further supported by our finding that several clusters are enriched for either genes with increased or decreased fitness. At the same time, our data demonstrates that the compensatory evolution of one module of the polarity pathway affects gene fitness in many other modules, including those that have no clear relation to cell polarization. These findings complement results from an earlier study by Harcombe et al. (74), which showed that gene loss can be compensated by mutations in functionally unrelated genes. Our results further demonstrate that, in addition to the compensatory mutations themselves, gene fitness across multiple cellular processes can change during compensatory evolution. It is conceivable that these changes in gene fitness will affect the further evolution of these modules. We therefore conclude that functional modules do not necessarily evolve independently of each other and that evolvability does not require modularity.

From our set of genes with a differential fitness, we were able to identify the bud scar, the polarisome and actin patches as the major structures that contribute to the latent pathway for polarity. While earlier studies indicated that actin based transport can drive polarization of Cdc42 (31), their results were criticized to be largely caused by the unintended activation of the stress response pathway. Our data provides new evidence that actin based transport can contribute to cell polarization, but its activity is most likely too weak to achieve polarity establishment on its own. The bud scar could provide an additional activation mechanism for Cdc42 that, in conjunction with actin recycling, is able to maintain a polarized distribution of Cdc42. Indeed, the bud-scar associated GTPase Rsr1 is capable of recruiting Cdc24, the GEF for Cdc42, to the bud site (75). This recruitment would cause a locally increased rate of Cdc42 activation near the bud scar, which, combined with a possible saturation of Cdc42 GAPs caused by the loss of Bem3, could drive polarity establishment. While such a mechanism largely agrees with the finding obtained by the model of Brauns et al. (39), a key difference is that we do predict that Cdc24 is polarized prior to budding in the polarity mutant. In addition, we propose that this latent polarity pathway may also depend on protein interactions that have not been observed in wild-type cells. For example, it is unclear why Boi1, which depends on Bem1 to interact with Cdc24, has an increased importance in the polarity mutant. The same goes for several other genes that are too numerous too discuss individually. Microscopy, *in vitro* experiments and molecular models will likely be needed to obtain a complete picture of the latent polarity pathway.

We found that the number of genes that increase fitness and those that decrease fitness after compensatory evolution is nearly equal. Thus, gene loss does not appear to necessarily decrease mutational robustness, but rather cause a restructuring of the gene-fitness relationship. While it is somewhat counter-intuitive that gene loss can lead to the increased dispensability of other genes, a mechanistic explanation has been given for this phenomenon. Specifically, if the loss of a protein fully inactivates an associated pathway, the additional loss of other proteins on the same pathway does not incur additional fitness costs (76, 77). In this way, the loss of a protein can appear to buffer mutations in other proteins.

However, this explanation does not hold for the case where gene deletions lead to the functional restoration of the perturbed pathway. Similarly, the scale at which we see changes in gene fitness occur and the diversity of cellular processes that are affected suggests an alternative mechanism. How then, can the loss of a small number of genes cause changes in the gene-fitness relation of such a large part of the genome? Although our results do not provide direct evidence, we propose that gene loss can induce widespread rewiring of the physical interaction network based on the following reasoning. The fitness effect resulting from the loss of a gene is related to the physiological role of the protein encoded by that gene. However, it is increasingly recognized that a gene does not encode for a protein with a static structure, but rather for structural ensembles (40, 78). Which structure is preferred depends on the environment, making protein function heavily context dependent (79). We speculate that in a similar manner, protein structure can be affected by the presence or absence of specific binding partners (42, 80). Loss of one binding partner could in that case cause a conformational change that affects its affinity for other possible binding partners. In this way, the deletion of one protein can initiate a cascade that induces large-scale rewiring of the protein interaction network.

In conclusion, we show that modularity plays a minor role during evolution. Instead, compensatory evolution induces genome-wide rearrangements in interaction networks that cross module boundaries and which might be mediated by protein promiscuity. Because these genome-wide changes affect the adaptive potential of many different cellular processes, this could explain how gene loss can promote evolutionary novelty (81–86) and allow populations to escape fitness peaks (82). Despite the minimal role of modularity, some hope for making evolutionary predictions might be drawn from our finding that genes related to the same biological process occasionally share the same sign in fitness change after compensatory evolution. Our work also makes a case for the importance of protein ensembles and weak interactions in shaping adaptive pathways. Interactions that are weak in non-perturbed conditions could be driving the re-establishment of biological pathways in response to perturbations. Mapping these weak, non-specific interactions that are frequently perceived as irrelevant will be instrumental to bridge the gap between biological network structure and the evolutionary potential of biological processes.

## 4. Methods

### A. Strains

All strains used in this study are from the W303 genetic background. The cas9 cassette was obtained from plasmid p414-TEF1p-cas9-CYC1t (87) and fused to the up- and downstream genomic sequences of the *HO*-locus and the *ScURA3* marker using overlap-extension PCR. The resulting genetic construct was transformed into the wild-type strain and the polarity mutant using according to a lithium-acetate transformation protocol (88). Correct integration was verified with colony PCR. Endogenous expression of Cas9 from the *HO*-locus had no significant effects on growth. Strains were made compatible with SATAY by removing the *ScURA3* marker and the endogenous *ADE2* locus using the CRISPR/Cas9 system according to the double guide-RNA (gRNA) method of Mans et al. (89). gRNA sequences targeting the *ScURA3* marker and *ADE2* locus were designed using the online toolbox CHOPCHOP (90). The repair fragment for removal of the *ScURA3* marker was constructed by PCR amplification of the genomic sequence upstream and downstream of the *ScURA3* marker and fusing the two fragments using overlap-extension PCR. Similarly, the repair fragment for removal of the *ADE2* locus was constructed by fusing the up- and downstream genomic sequences with overlap-extension PCR. Removal of *ScURA3* marker and the *ADE2* was verified by colony PCR using primers 5 and 6 and primers 7 and 8, respectively. Correct clones were cured from their gRNA plasmids by inoculating them in non-selective medium (YPD) and growing them until saturation (∼ 1.5 days) at 30 °C. After saturation was reached, the cultures were plated to single colonies on YPD agar plates and screened for the loss of growth on media selecting for the marker of the gRNA plasmid. Strains were stored at −80 °C as frozen stocks in 40% (v/v) glycerol. The double gRNA plasmids that were used as a template were a kind gift from Pascale Daran-Lapujade.

### B. Media

Standard culturing and growth assays were performed in YPD (10g/L Yeast extract, 20 g/L Peptone, 20 g/L dextrose), SC (6.9 g/L Yeast nitrogen base, 0.75 g/L Complete supplement mixture, 20 g/L dextrose). For *ade−* strains, standard growth media was supplemented with 20 mg/L adenine just before incubation. Liquid media for the preculture and induction steps of SATAY were prepared according to the recipe in table S1. After preparation, the media was filter sterilized using Rapid-Flow Sterile Disposable Filter Units (Nalgene) and stored at 4°C until use. Liquid media for the reseed step of SATAY was prepared by autoclaving 2.6 L of MiliQ water in a 5 L flask. 400 mL of a 7.5X concentrated solution of the nutrients was prepared separately and filter sterilized. To prevent the degradation of media components, this concentrate was stored in the dark at 4°C until used. On the day of reseed, the concentrate was aseptically added to the 5 L flask containing 2.6 L of MiliQ water and mixed. Solid media was prepared by adding 20 g/L agar and 30 mM Tris-HCl (pH 7.0) to the liquid media recipe and autoclaving the mixture for 20 minutes at 121°C. 20 mg/L adenine was aseptically added after autoclaving, unless plates were intended to be selective for adenine auxotrophy.

### C. Transposon mutagenesis screens

#### C.1 Library generation

SATAY libraries were generated based on the procedure described by (91), which is a modification of the original protocol (92) to allow transposition in liquid media. *ade−* cells were transformed with plasmid pBK549 (91), which was a kind gift from Benoît Kornmann, according to a lithium acetate transformation protocol (88). To screen for clones transformed with the intact version of plasmid pBK549 (see (93) for details on the different species of pBK549), 12-24 colonies were picked from the transformation plate, re-streaked on fresh SD-ADE and SD-URA plates and incubated for 3 days at 30°C. For clones that showed full growth on SD-URA plates while producing a small number of colonies on SD-ADE plates, cells were scraped from the SD-URA plate and used to inoculate 25 mL of preculture media at an OD600 of 0.20-0.28. Precultures were grown on an orbital platform shaker at 160 rpm, 30 °C until the OD600 was between 5-7 (∼20h). The saturated precultures were used to inoculate 200 mL of induction media at an OD600 of 0.10-0.27 and grown for 52 hours to allow transposition to occur. The efficiency of transposition was monitored by plating samples of the liquid induction cultures on SD-ADE at T=0 and T=52 hours and scoring the number of colonies on these plates after 3 days of incubation at 30 °C. After 52 hours of induction, the resulting transposon mutagenesis libraries were reseeded in 3 liters of reseed media at an OD600 of 0.21-0.26. Typically, this meant that around 7 million transposon mutants were reseeded per library. Reseeded libraries were grown for 92 hours at 140 rpm, 30 °C. At the end of reseed, cells were harvested by centrifugation of the reseed cultures at 5000 xg for 30 minutes. Cell pellets were stored at −20 °C.

#### C.2 Genomic DNA extraction

A 500 mg frozen pellet was resuspended in 500 µL cell breaking buffer^∗^ and distributed into 280 µL aliquots. 300 µL of 0.4-0.6 mm glass beads (Sigma-Aldrich, G8772) and 200 µL of Phenol:Chloroform:isoamyl alcohol 25:24:1 (Sigma-Aldrich, P2069) were added to each aliquot and cells were lysed by vortexing the samples with a Vortex Genie 2 at maximum speed at 4°C for 10 minutes. 200 µL of TE buffer was added to each lysate, after which the samples were centrifuged at 16100x g, 4°C for 5 minutes. After centrifugation, the upper layer (∼400 µL) was transferred to a clean eppendorf tube. 2.5 volumes of 100% absolute ethanol was added to each sample and mixed by inversion to precipitate the genomic DNA. After precipitation, the DNA was pelleted by centrifugation at 16100x g, 20°C for 5 minutes. The supernatant was removed and the DNA pellet was resuspended in 200 µL of 250 µg/ml RNAse A solution (qiagen, Cat. No. 19101). the resuspended DNA pellets were incubated at 55°C for 15 minutes to allow digestion of the RNA. After digestion, 20 µL 3M, pH 5.2 sodium acetate (Merck) and 550 µL 100% absolute ethanol was added to each sample and mix by inversion. DNA was pelleted by centrifugation at 16100x g, 20°C for 5 minutes. Pellets were washed with 70% absolute ethanol and dried at 37°C for 10 minutes or until all ethanol had evaporated. The dried pellets were resuspended in a total volume of 100 µL MiliQ water and the concentration of the genomic DNA samples was quantified on a 0.6% agarose gel using the Eurogentec Smartladder 200bp-10kb as a reference. Prepared DNA samples were stored at −20°C or 4°C until used.

#### C.3 Library sequencing

To prepare genomic DNA samples for sequencing, 2×2 µg of DNA from each sample were transferred to non-stick microcentrifuge tubes and digested with 50 units of DpnII and NlaIII in a total volume of 50 µL for 17 hours at 37°C. After digestion, the restriction enzymes were heat-inactivated by incubating the samples at 65°C for 20 minutes. Digestion results were qualitatively assessed by visualization on a 1% agarose gel stained with Sybr-Safe. Successfully digested DNA samples were circularized in the same tube using 25 Weiss units of T4 DNA ligase (Thermo Scientific, Catalog #EL0011) at 22°C for 6 hours in a total volume of 400 µL. After ligation, the circularized DNA was precipitated using 1ml 100% absolute ethanol, 20 µL 3M, pH 5.2 sodium acetate (Merck) and 5 µg linear acrylamide (invitrogen, AM9520) as a carrier. DNA was precipitated for at least 2 days at −20°C. Precipitated DNA was pelleted by centrifugation for 20 minutes at 16100x g at 4°C and washed with 1 ml of 70% ethanol. After washing, the DNA was re-pelleted by centrifugation for 20 minutes at 16100x g at 20°C, the supernatant was removed and pellets were dried for 10 minutes a 37°C. Each dried pellet was resuspended in water and used as a template for 20 PCR reactions of 50 µL. Transposon-genome junctions were amplified using the barcoded primers 1 and 2 (table S2) for DpnII digested DNA or primers 3 and 4 (table S2) for NlaIII digested DNA on a thermal cycler (Bio-Rad C1000 Touch). PCR amplified samples were purified using the NucleoSpin Gel and PCR cleanup kit (Macherey-Nagel) and quantified on the NanoDrop 2000 spectrophotometer (Thermo Scientific). For each sample, equal ratios (w/w) of DpnII and NlaIII digested DNA were pooled. Library preparation and sample sequencing were performed by Novogene (UK) Company Limited. Sequencing libraries were prepared with the NEBNext Ultra II DNA Library Prep Kit, omitting the size selection and PCR enrichment steps. DNA libraries were sequenced on the Illumina NovaSeq 6000 platform using Paired-End (PE) sequencing with a read length of 150 bp.

#### C.4 Sequence alignment

FASTQ files obtained from the NovaSeq 6000 platform were demultiplexed into DpnII and NlaIII digested DNA samples based on the barcodes introduced during PCR amplification. Read pairs with non-matching barcodes were discarded. After demultiplexing, the forward read of each read pair was selected and the sequences upstream of primer 688_minidsSEQ1210 (43) and downstream of the DpnII (GATC) or NlaIII (CATG) restriction site were trimmed. All demultiplexing and trimming steps were performed with BBduk (94) integrated into a home-written pipeline written in Bash. After trimming, the forward reads were aligned to the S288C reference genome (version R64-2-1_20150113) with the Transposonmapper pipeline ((95) version v1.1.4) using the following settings:

- Data type: ‘Single-end’
- Trimming software: ‘donottrim’
- Alignment settings: ‘-t 1 -v 2’

### D Volcano plots

Read count distributions were corrected for spikes using the Beta-Geometric correction method and subsequently normalized for differences in transposon density and sequencing depth with the median of ratios normalization (see section A). P-values were generated using the unequal variance independent t-test available from the SciPy library in Python (96) and corrected for multiple hypothesis testing with the Benjamini-Hochberg procedure implemented in the TRANSIT software tool ((97), version 3.2.6). Genes were considered to be essential if they have been annotated as such by the Saccharomyces Genome Database ((98), accessed on 03/17/2006).

### E Physical interaction network analysis

Because the correlation between evolutionary dynamics and PPI topology depends on network quality, we only included interactions from the Multi-Validated dataset. The Multi-Validated (MV) dataset of physical interactions (release BIOGRID-4.4.214) was downloaded from the BioGRID in Tab 3.0 format, which contains all physical interactions that have been validated by at least two different experimental systems or publication sources. These experimental systems include high-throughput *in vitro* screens, and the network therefore contains a mixture of *in vitro* and *in vivo* validated interactions. Moreover, it should be noted that the constructed PPI is a static representation of a dynamic network structure. The original set of physical interactions was filtered to obtain only those that correspond to interactions between proteins of *S. cerevisiae*. Interactions inferred from Affinity Capture-RNA or Protein-RNA were excluded from the dataset as these interactions do not represent direct physical interactions between proteins. Network visualizations were made with Cytoscape ((99), version 3.9.1). Network centralities (degree, betweenness and closeness) were calculated using the NetworkX package in Python ((100), version 2.8.4).

### F Functional interaction network analysis

Functional associations between gene products were retrieved from the STRING database (51) through the stringApp plugin ((52), version 1.7.1) of Cytoscape ((99), version 3.9.1). The confidence threshold of the imported interactions from STRING was set to medium (confidence level ≤ 0.40). Markov Clustering (MCL) was performed with the clusterMaker2 (version 2.2) plugin of Cytoscape using the confidence level as edge weight and setting the granularity parameter to 2.5.

## 5. Acknowledgements

We thank Agnès Michel and Benoît Kornmann for providing the plasmid pBK549 containing the MiniDS transposon system, their help and guidance when setting up the SATAY assays and insightful discussions. We are grateful to Melanie Wijsman and Nicole Bennis for providing us with the plasmids for CRISPR/Cas9 gene editing and their support during troubleshooting. We also thank Leila Iñigo de la Cruz and Gregory van Beek for establishing and maintaining the SATAY data analysis pipeline and their advice regarding data management. L.L. and E.K. gratefully acknowledge funding from the European Research Council under the European Union’s Horizon 2020 research and innovation programme (grant agreement 758132).

## 6. Supplement

### A. Read count correction and normalization procedure

The benefit of including read counts in our metric to identify changes in the fitness effect of gene disruptions across genetic backgrounds is that it allows for a more sensitive analysis than those based on insertion counts. For example, a gene may have a negligible difference in insertion counts between two genetic backgrounds if the change in gene disruption tolerance is not so drastic that it causes a large number of mutants to fall below the detection level in one of the two backgrounds. However, if a significant difference in gene disruption tolerance truly exists, we would expect that the read counts associated with each insertion site (which reflect mutant abundance) will be lower in one of the two strains. Thus, extending the analysis from a binary assessment of the occupancy of insertion sites to a gradual scale based on read counts can uncover differences that would otherwise remain undetected. However, a drawback is that read counts tend to be more variable between replicate experiments than insertion counts, often owing to read count spikes caused by technical and biological artifacts (101). This increased noise level can hinder the desired increase in sensitivity of the analysis. In addition, spikes in the read counts can significantly skew the read count distribution. This means that the small subset of insertion sites with an artificially high read count will strongly affect methods that aim to correct differences in sequencing depth between libraries by a linear transformation of the total read count.

To address these issues, we implemented the Beta-Geometric correction (BGC) method developed by DeJesus and Ioerger (102). This method is based on the observation that read count distributions obtained from transposon insertion sequencing (TIS) experiments resemble a geometric distribution where most insertion sites have a low read count, while only few sites contain many reads. During the BGC correction procedure, the quantiles of the empirical read count distribution are adjusted to match the quantiles of a fitted ‘ideal’ geometric distribution. To allow for greater flexibility of the fit, this ideal distribution is implemented as a mixture of geometric distributions with a Beta prior on the probability parameter. In practice, this procedure corrects for the ‘skew’ in the empirical read count distribution by suppressing spikes while inflating the insertion sites that have a low read count. The results in figure S1 show that the application of this procedure greatly improves the correlation between replicate datasets (figure S1b) compared to uncorrected datasets (figure S1a). Lastly, after applying the BGC correction, we normalized for differences in sequencing depth and library complexity across datasets using the median of ratios normalization (103, 104). In summary, the geometric mean across all samples was calculated for each gene:

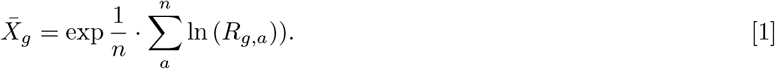

Where 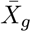 is the geometric mean of the read counts mapping to gene *g, n* is the total number of datasets and *R*_*g,a*_ is the number of reads that map to gene *g* in dataset *a*. Next, the ratio of the total read count to the geometric mean 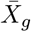 was determined for each sample and the sample-specific normalization factor was taken to be the median of these ratios across all genes:

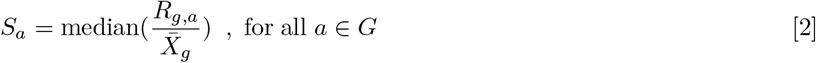

With *S*_*a*_ the normalization factor for dataset *a* and *G* is the complete set of annotated genes used in this chapter. Finally, the read counts were linearly scaled by the normalization factor *S*_*a*_:

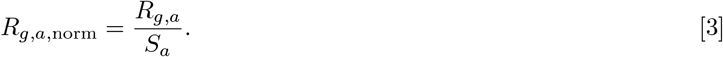

### B. Supplemental figures

**Fig. S1.**
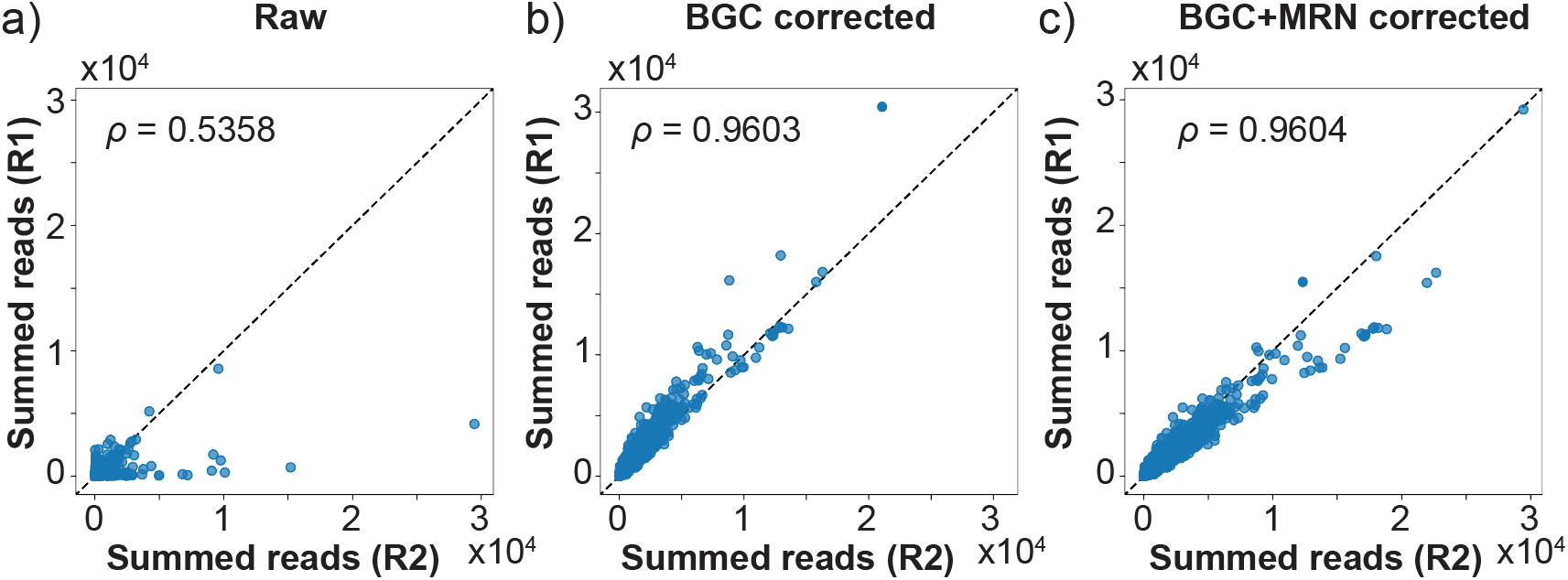
Normalization procedure increases correlation in the summed read counts per gene between replicate experiments. In order to improve the correlation between replicate datasets derived from the same genetic background, the datasets are first aligned with their best fitting Beta-Geometric distribution (panel (a) to panel (b)) to correct for spikes in read counts, followed by a median of ratios normalization to account for differences in sequencing depth (panel (b) to panel (c)). In each panel, the value of Pearson’s correlation coefficient (*ρ*) is shown.

**Fig. S2.**
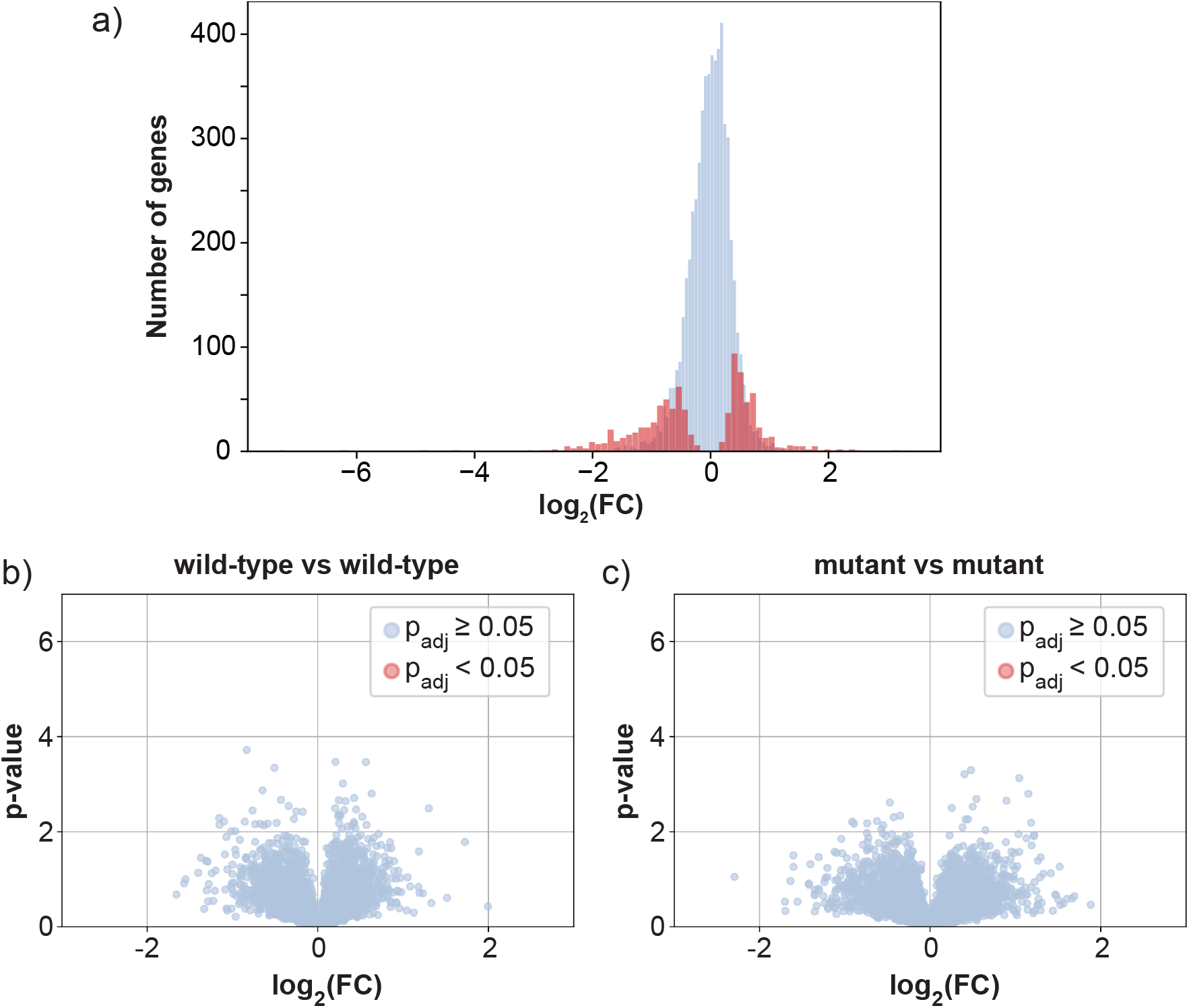
Validity of the geneset that is identified as having differential fitness between the wild-type strain and the polarity mutant. (**a**) Significant genes have a larger effect size. Histogram of the log_2_-fold changes shown in figure 2. The plot shows that genes that are flagged to have a statistically significant difference between the two genetic backgrounds (red bars) typically occurs for larger fold-changes. (**b-c**) Comparing transposon mutagenesis libraries obtained from the same genetic background yields no significant differences in gene fitness. Volcano plots are shown for comparisons between wild-type and mutant datasets. The replicate datasets of each genetic background (6 in total) were split and compared 3 vs. 3. For both our wild-type strain and from our polarity mutant, no false positives are found at a significance threshold of p_adj_ *<* 0.05.

**Fig. S3.**
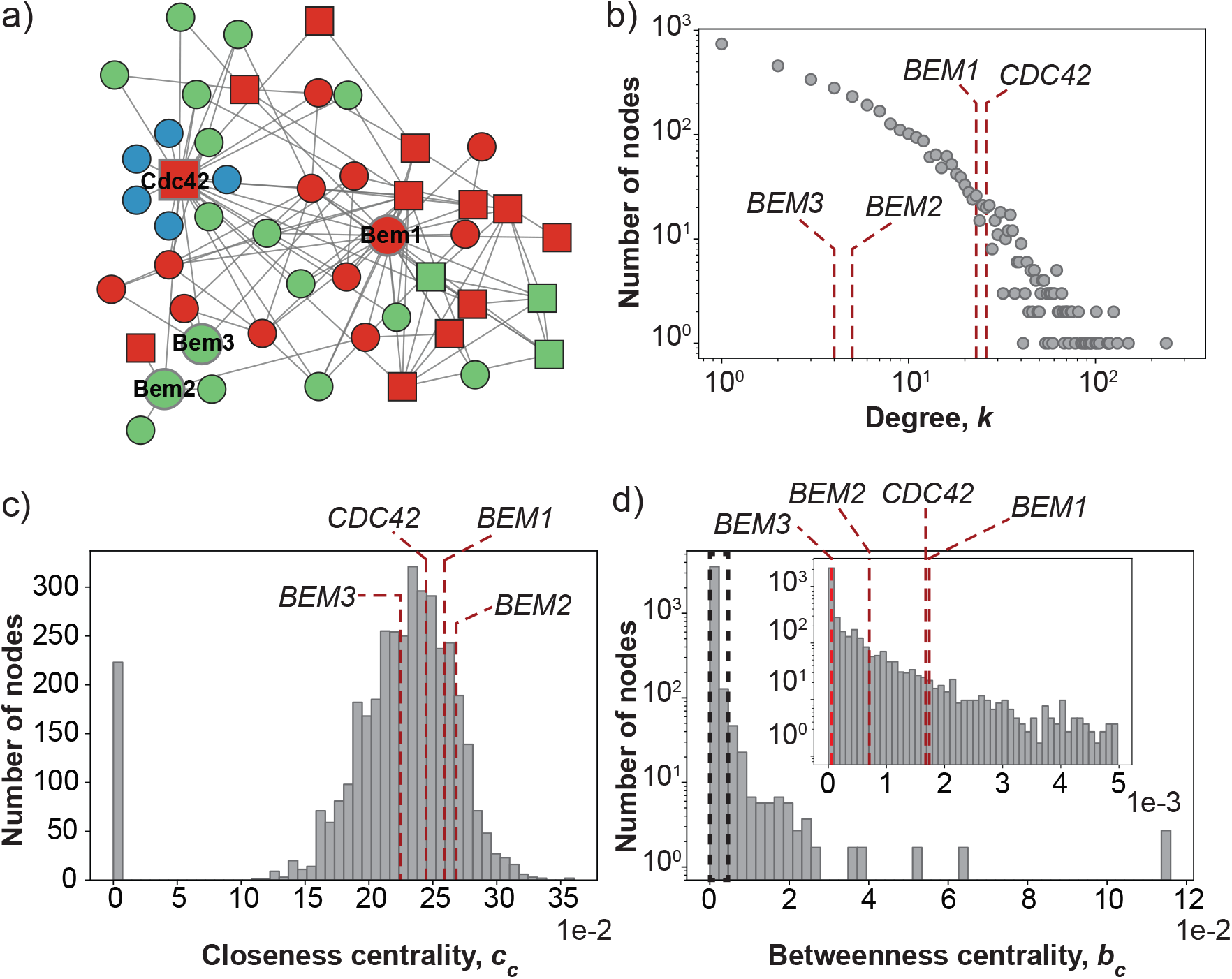
Properties of the constructed protein-protein interaction network. (**a**) Sub-graph of of the protein-protein interaction network for Cdc42, Bem1, Bem2 and Bem3 and their first neighbours. Nodes are colored according to their degree *k* in the complete PPI network (figure 3a). Blue: *k ≤* 3, green: 4 *≤ k ≤* 10, red: *k >* 10. Proteins are essential according to the SGD database are shown as squares, non-essential proteins are shown as circles. (**b**) The degree centrality of the nodes in the complete PPI network. The degree distribution shows the typical sub-linearity of PPI networks in biology when plotted on a log-log scale. (**c**) The closeness centrality distribution of the PPI network. (**d**) the betweenness centrality distribution of the PPI network. In the panels **c**-**d**, the degree values are indicated for two polarity proteins that have a strong negative effect on fitness when deleted (Bem1 and Cdc42) and for two polarity proteins that have a moderate to weak negative effect (Bem3 and Bem2). With respect to degree and betweenness, proteins with a similar gene fitness lie in proximity of each other on the distributions. For the closeness distribution, we find no relation with gene fitness.

**Fig. S4.**
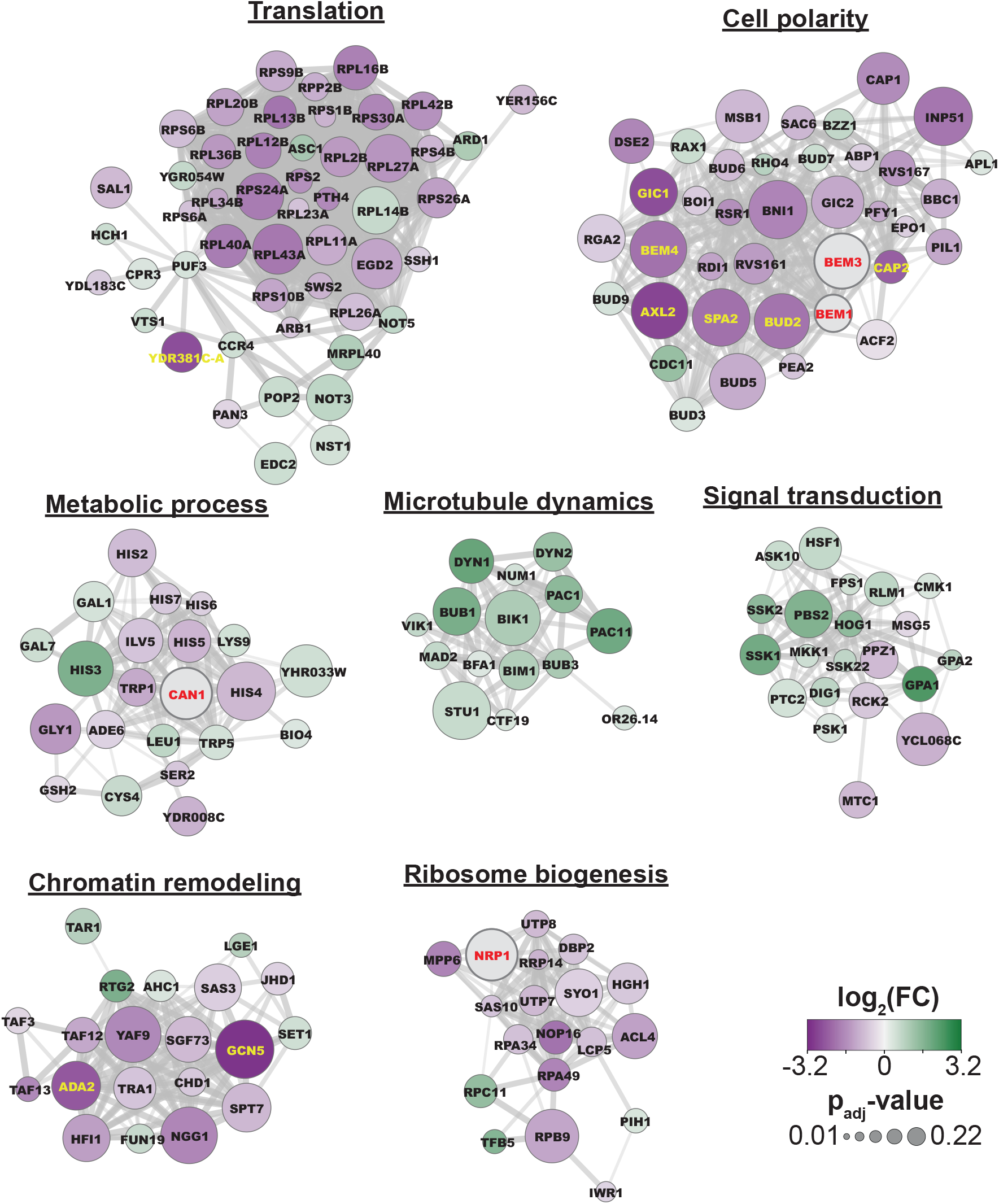
Zoom-in of the seven largest clusters identified in our functional association network by the Markov clustering algorithm. The biological process gene ontology enrichment is shown above each cluster.

### C. Supplemental tables

**Table S1.** Liquid media used for SATAY library generation. YNB: Yeast Nitrogen Base. CSM: Complete Supplement Mixture. Ura: Uracil. Ade: Adenine.

| Step | Media | Components |
| --- | --- | --- |
| Preculture | SD-Ura+0.2% Glucose<br>+2% Raffinose | YNB w/o Amino Acids (6.8 g/L)<br>CSM -Ura (0.77 g/L)<br>Glucose (2 g/L)<br>Raffinose (20 g/L)<br>Adenine (20 mg/L) |
| Induction | SD-Ura+2% Galactose | YNB w/o Amino Acids (6.8 g/L)<br>CSM -Ura (0.77 g/L)<br>Galactose (20 g/L)<br>Adenine (20 mg/L) |
| Reseed | SD-Ade+2% Glucose | YNB w/o Amino Acids (6.8 g/L)<br>CSM -Ade (0.78 g/L)<br>Glucose (20 g/L) |

**Table S2.**
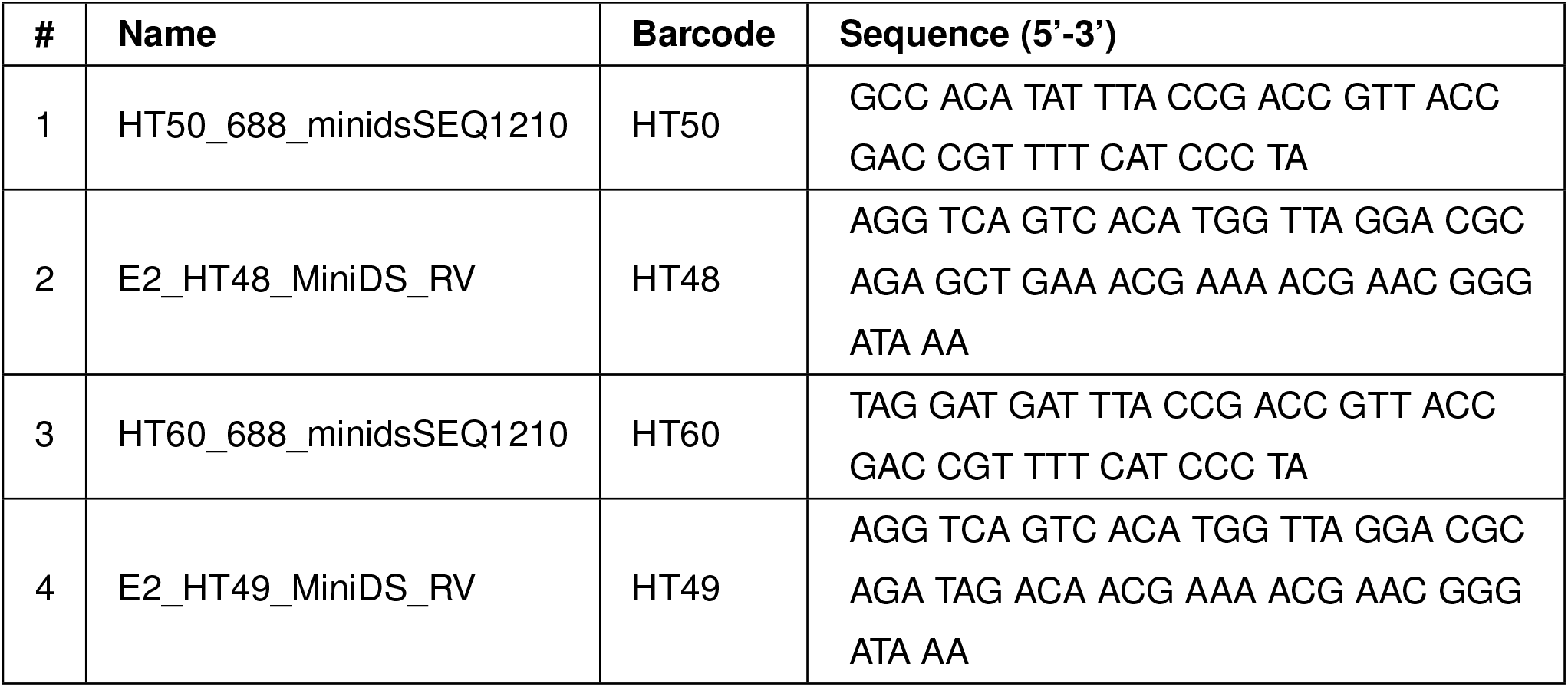
List of used primers.

## Footnotes

∗ 2% Triton X-100, 1% SDS, 100 mM NaCl, 100 mM Tris-HCl pH8.0, 1 mM EDTA

